# C-terminal HA tags compromise function and exacerbate phenotypes of *Saccharomyces cerevisiae* Bloom’s helicase homolog Sgs1 SUMOylation-associated mutants

**DOI:** 10.1101/2020.04.23.058578

**Authors:** Matan Cohen, Michael Lichten

## Abstract

The Sgs1 helicase and Top3-Rmi1 decatenase form a complex that affects homologous recombination outcomes during the mitotic cell cycle and during meiosis. Previous studies have reported that Sgs1-Top3-Rmi1 function is regulated by SUMOylation that is catalyzed by the Smc5-Smc6-Mms21 complex. These studies used strains in which *SGS1* was C-terminally tagged with three or six copies of a human influenza hemagglutinin-derived epitope tag (3HA and 6HA). They identified *SGS1* mutants that affect its SUMOylation, which we will refer to as *SGS1* SUMO-site mutants. In previous work, these mutants showed phenotypes consistent with substantial loss of Sgs1-Top3-Rmi1 function during the mitotic cell cycle. We find that the reported phenotypes are largely due to the presence of the HA epitope tags. Untagged *SGS1* SUMO-site mutants show either wild-type or weak hypomorphic phenotypes, depending on the assay. These phenotypes are exacerbated by both 6HA and 3HA epitope tags in two different *S. cerevisiae* strain backgrounds. Importantly, a C-terminal 6HA tag confers strong hypomorphic or null phenotypes on an otherwise wild-type Sgs1 protein. Taken together, these results suggest that the HA epitope tags used in previous studies seriously compromise Sgs1 function. Furthermore, they raise the possibilities either that sufficient SUMOylation of the Sgs1-Top3-Rmi1 complex might still occur in the SUMO-site mutants isolated, or that Smc5-Smc6-Mms21-mediated SUMOylation plays a minor role in the regulation of Sgs1-Top3-Rmi1 during recombination.

## Introduction

DNA double strand breaks (DSBs) present a major threat to genome integrity and, if repaired incorrectly, can lead to a loss of genetic information. Cells have developed multiple mechanisms to repair these lesions, and the safest repair process is homologous recombination (HR). During HR, a broken DNA strand invades a homologous chromosome and uses it as a repair template. Repair can occur with or without exchange of chromosome arms, producing crossover (CO) or noncrossover (NCO) recombinants, respectively (Kowalczykowski 2015). During the mitotic cell cycle, cells repress CO formation and favor the NCO outcome (LaRocque *et al*. 2011; Symington *et al*. 2014). During meiosis, cells use HR to promote homologous chromosome alignment and segregation during the first nuclear division. This requires the formation of regulated crossover products by only a subset of the initiating DSBs (Hunter 2015; Zickler and Kleckner 2015; Lam *et al*. 2017). A group of meiosis-specific and biochemically diverse factors, referred to collectively as the ZMM proteins, collaborate to stabilize strand invasion intermediates and to promote formation of double Holliday junctions (dHJs). ZMM-promoted dHJs are resolved predominantly as crossovers by the action of the Mlh1-Mlh3-Exo1 (MutLγ) complex (Fung *et al*. 2004; Snowden *et al*. 2004; Lynn *et al*. 2007; De Muyt *et al*. 2012; Zakharyevich *et al*. 2012; Hunter 2015).

The Sgs1-Top3-Rmi1 (STR) helicase-decatenase complex and its homologs are central regulators of recombination product formation during both the mitotic and meiotic cell cycles (Ira *et al*. 2003; Jessop *et al*. 2006; Oh *et al*. 2007; Jessop and Lichten 2008; Oh *et al*. 2008; Larocque *et al*. 2011; De Muyt *et al*. 2012; Zakharyevich *et al*. 2012; Hunter 2015; Kaur *et al*. 2015; Tang *et al*. 2015). STR and homologs are thought to promote NCO formation by unwinding strand invasion intermediates in a process known as synthesis dependent strand annealing (SDSA)(Cejka and Kowalczykowski 2010; Fasching *et al*. 2015). STR and its homologs can also disassemble dHJs and form NCOs in a process known as dissolution (Wu *et al*. 2005; Cejka and Kowalczykowski 2010; Dayani *et al*. 2011; Kaur *et al*. 2019). In addition, the Top3-Rmi1 subcomplex has an Sgs1-independent role in the resolution of recombination intermediates (Kaur *et al*. 2015; Tang *et al*. 2015). During meiosis, the D-loop disassembly activity of the STR complex is hypothesized to lead to recycling of early strand invasion intermediates, which can promote NCO formation or promote recombination intermediate stabilization by the ZMM proteins and subsequent resolution as COs (Jessop *et al*. 2006; De Muyt *et al*. 2012; Zakharyevich *et al*. 2012; Hatkevich and Sekelsky 2017).

Two recent studies have proposed a mechanism for STR complex activity regulation by the Smc5-Smc6-Mms21 complex (Bermudez-Lopez *et al*. 2016; Bonner *et al*. 2016). The Smc5-Smc6-Mms21 complex is a member of the SMC (Structural Maintenance of Chromosomes) family with structural similarities to cohesin and condensin, and is important in chromosome transactions such as DNA replication and repair. The Smc5-Smc6-Mms21 complex is unique among SMC complexes because it contains an essential subunit, Nse2/Mms21 (referred to as Mms21 here), with an SP-RING domain in its C-terminus that contains E3 SUMO ligase activity (Andrews *et al*. 2005; Potts and Yu 2005; Zhao and Blobel 2005; Aragon 2018). In budding yeast, mutants lacking this E3 SUMO ligase activity are viable but are highly sensitive to DNA damage (Zhao and Blobel 2005). The two studies of SUMO-mediated STR regulation referred to above (Bermudez-Lopez *et al*. 2016; Bonner *et al*. 2016) suggested that DNA lesions promote Mms21-mediated SUMOylation of Smc5-Smc6-Mms21 components, which then act as a platform to recruit STR through Sgs1’s <u>S</u>UMO <u>I</u>nteraction <u>M</u>otifs (SIMs). This Sgs1-Smc5 interaction is then suggested to result in Mms21-mediated modification of STR components, which in turn promotes STR activity during homologous recombination (Bermudez-Lopez *et al*. 2016; Bonner *et al*. 2016). In these studies, which used either a 6HA (Bermudez-Lopez *et al*. 2016) or a 3HA (Bonner *et al*. 2016) epitope tag at the Sgs1 C-terminus, Sgs1 was found to be SUMOylated at 6 lysines, with lysing 621 (K621) being the major site. Lysine to arginine mutants at either K621 or at all 6 lysines substantially reduced MMS-induced SUMOylation without affecting Sgs1-Smc5 interaction (Bermudez-Lopez *et al*. 2016; Bonner *et al*. 2016). The two groups mutated different sets of residues in Sgs1 SIMs, but both sets of mutants blocked Sgs1-Smc5 interaction while leaving Sgs1-Top3 interaction intact (Bermudez-lopez *et al*. 2016; Bonner *et al*. 2016).

Work in human cells has also revealed a role for SUMOylation of the Sgs1 homolog, BLM, in rescuing of stalled replication forks (Ouyang *et al*. 2009). This study proposed that SUMOylation of BLM relieves the inhibition of RAD51 at a stalled replication fork, and allows for homologous recombination to proceed. Furthermore, this activity was found to be dependent on the NSMCE2 protein, the human homolog of Mms21 (Pond *et al*. 2019). These studies suggest that SUMOylation of BLM/Sgs1 may be evolutionarily conserved. However, the nature of the contribution of this modification to BLM/Sgs1 activity and the different contexts in which this modification is required are not well understood.

The SUMO ligase activity of Mms21 is required for the Smc5-Smc6-Mms21 complex’s role in destabilizing aberrant intermediates formed during the early stages of meiotic recombination (Xaver *et al*. 2013), and many of the meiotic phenotypes of Smc5-Smc6-Mms21 mutants closely resemble those of mutants lacking STR activities (Jessop *et al*. 2006; Oh *et al*. 2007; Jessop and Lichten 2008; Oh *et al*. 2008; De Muyt *et al*. 2012; Copsey *et al*. 2013; Lilienthal *et al*. 2013; Xaver *et al*. 2013; Kaur *et al*. 2015; Tang *et al*. 2015). However, it is not known if the requirement for Mms21 SUMO E3 ligase activity reflects SUMOylation of the STR complex or of other proteins involved in recombination.

We wished to test the hypothesis that the defects of *mms21* mutants can be attributed to an absence of STR SUMO modification. We reasoned that mutations in Sgs1 that block its SUMOylation should not alter STR recruitment by Smc5-Smc6-Mms21 or modification of Top3 and Rmi1, whereas mutations in Sgs1 SIMs would prevent STR recruitment and thus abolish modification of all of the STR components. Therefore, we recreated lysine to arginine mutants at either the major SUMOylation site (*sgs1-K621R*) or all 6 lysines previously identified as being SUMOylated (hereafter called *sgs1-6KR*, referred to by Bermudez-Lopez *et al*. 2016 as *sgs1-3KR*), and also mutated the residues within the SIMs previously mutated by the Zhao (Bonner *et al*. 2016) and Aragon (Bermudez-Lopez *et al*. 2016) groups (referred to as *sgs1-ZSIM* and *sgs1-ASIM*, respectively; see Fig. 1A for details of all mutants). We integrated these mutants, which we will refer to collectively as *SGS1* SUMO-site mutants, at the endogenous *SGS1* locus, either with or without a C-terminal epitope tag, and examined the resulting strains for both mitotic and meiotic STR function.

**Figure 1.**
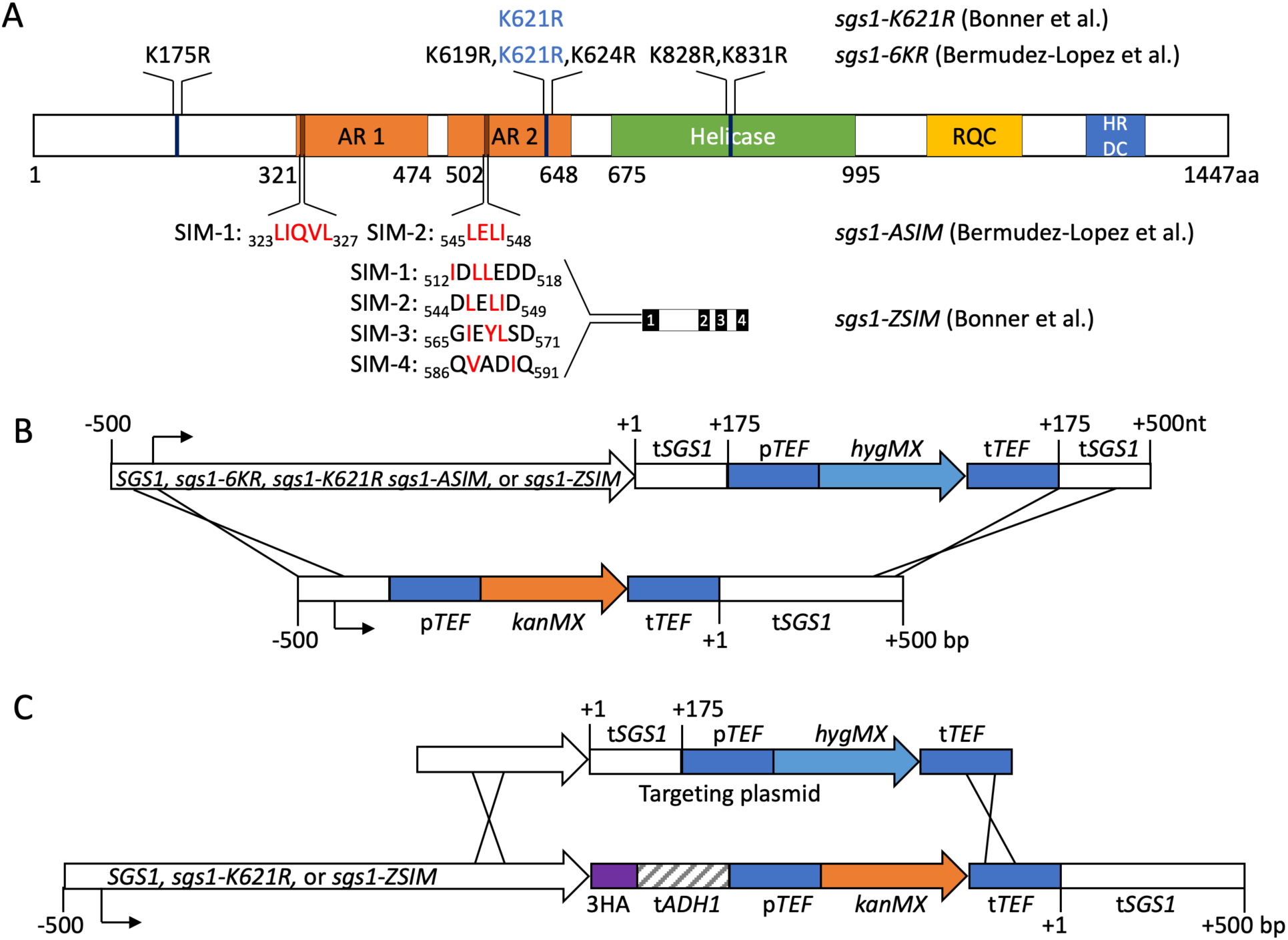
*SGS1* SUMO-site mutants and allelic replacement strategy. A) Map of the Sgs1 protein and the SUMO-site mutants used in this study. Top–K to R mutations used in Bonner *et al*. (2016) and Bermudez-Lopez *et al*. (2016); mutation in blue is common to both studies. *sgs1-6KR* is called *sgs1-3KR* in Bermudez-Lopez *et al*.(2016). Bottom—SUMO-interaction module (SIM) mutants. Residues in red were replaced by alanine in the indicated report. Sgs1 domains are: AR1, AR2—acidic regions 1 and 2; RQC—RecQ C-terminal domain; HRDC—helicase-and-RNAseD-like-C-terminal domain. Numbers indicate amino acid residues. B) Allelic replacement strategy used to insert *SGS1* SUMO-site mutants in the genome of SK1 strains. Plasmids containing mutants were digested to release the indicated fragment, which used homologous sequences up- and downstream of the *SGS1* ORF to direct integration. Numbers indicate nucleotides relative to the *SGS1* ORF start (negative numbers) and the *SGS1* ORF end (positive numbers). Promoters and terminators are indicated by the prefixes “p” and “t,” respectively. Construct segments are not drawn to scale. C) Strategy used to remove the 3xHA tag from *SGS1* mutants in W303 strains from Bonner *et al*. (2016). The indicated PCR fragment, which contains *SGS1* sequences 3’ of mutant-containing sequences, was used to replace sequences downstream of *SGS1*. Numbers are as in (B). Construct segments are not drawn to scale.

Our findings suggest that the previously reported phenotypes of the *SGS1* SUMO-site mutants are largely due to the presence of C-terminal HA epitope tags. In particular, a 6HA tag rendered an otherwise wildtype *SGS1* gene nonfunctional, and SUMO-site mutant strains lacking an epitope tag showed only mildly hypomorphic phenotypes with regards to MMS sensitivity and synthetic interactions with *slx4* mutation. We also find that untagged SUMO-site mutant strains do not show the meiotic viability defects and the ZMM mutant bypass phenotypes displayed by *sgs1* loss-of-function mutants, nor do they display the synthetic nuclear division defects that *sgs1* loss-of-function mutants do when combined with *mms4* mutants. These results point to a role for the Sgs1 C-terminus in STR function. They also raise the possibilities that SUMOylation may not play an essential role in regulation of the STR complex, and that regulation of recombination by the E3 SUMO ligase function of Mms21 may operate through targets other than the STR complex.

## Materials and Methods

### Yeast Strains

Strains used in this study are *S. cerevisiae* of SK1 (Kane and Roth 1974) or W303 (Bonner *et al*. 2016) backgrounds (Table S1). Strains were constructed by transformation or genetic crosses. *SGS1* allelic replacements were made by transforming an *sgs1Δ* strain (deleted for all *SGS1* coding sequences) with an allelic replacement fragment that contains, in the following order: 500 nt immediately upstream of the *SGS1* start codon; the entire *SGS1* coding sequence; 175 nt immediately downstream of the *SGS1* stop codon; a *hygMX* cassette (Janke *et al*. 2004); and the next 361 bp downstream of *SGS1* (Fig. 1B). In some cases, a C-terminal tag was included. Mutant alleles were inserted into a plasmid with this fragment by Gibson assembly, and were confirmed by sequencing. The 3HA tag in the W303 strains was removed by transformation with a PCR product containing the last 890 bp of *SGS1* through TEF terminator sequences in the *hygMX* cassette from the wild-type allelic replacement plasmid (Fig. 1C). Yeast transformants were confirmed by Southern blotting to ensure proper integration, and PCR fragments from transformants were sequenced to confirm 3’ end and terminator structures.

### Sporulation and spore viability

Diploid strains were grown in pre-sporulation media and sporulated as previously described (Goyon and Lichten 1993; Boerner and Cha 2015). For spore viability analysis, tetrads were collected after 24 h of sporulation, and at least 70 tetrads per genotype were dissected. For experiments examining synthetic interactions, haploids were mated and then sporulated on solid media for 1-2 days before dissection.

### Cytology

Nuclear divisions were monitored by DAPI staining of cells from liquid sporulations as previously described (Kaur *et al*. 2018). At least 200 cells were scored per time point.

### MMS and HU sensitivity assays

Cells were grown overnight in YPD liquid media (Kaur *et al*. 2018). After adjusting cell concentrations to the same OD_600_, cells were diluted in a 10-fold series, spotted on YPD plates containing the indicated drug concentrations, incubated at 30° C for 2-3 days, and imaged.

### Statistical analysis

GraphPad Prism was used for comparisons of spore viability using Fisher’s exact test, applying the Bonferroni correction for multiple comparisons.

### Data Availability

All strains and plasmids are available upon request without restrictions. Underlying data for spore viability and meiotic progression are available in Supplementary File 1.

## Results

### Validation of the allelic replacement strategy

As part of a project investigating whether the meiotic defects of Smc5-Smc6-Mms21 complex mutants can be attributed to a loss of STR complex recruitment or modification, we analyzed several *SGS1* mutants that were characterized in two recent studies of Sgs1 SUMOylation (Fig. 1A). These two studies used *SGS1* alleles with C-terminal epitope tags and non-native terminators (Bermudez-Lopez *et al*. 2016; Bonner *et al*. 2016). To avoid potential complications of these non-native configurations, all mutant alleles used in our study, as well as a wild-type control, were generated as allelic replacements that retain the native *SGS1* promoter and terminator (Fig. 1B and Materials and Methods). *sgs1Δ* mutants are sensitive to DNA damaging agents, such as MMS and HU, and exhibit synthetic lethality with *slx4Δ*, a phenotype shared with *mms21* mutants that lack SUMOylation activity (Mullen *et al*. 2000; Mullen *et al*. 2001; Zhao and Blobel 2005; Xaver *et al*. 2013). The untagged wild-type allelic replacement was not synthetically lethal with *slx4Δ* (Fig. 2A), nor was it sensitive to MMS or hydroxyurea (HU) (Fig. 2C), indicating that this allelic replacement strategy did not abrogate *SGS1* function (Mullen *et al*. 2001; Ui *et al*. 2001). Using this approach, we recreated the *SGS1* SUMO-site mutants and set out to analyze whether these mutants recapitulate previously reported STR or Smc5-Smc6-Mms21 mutant phenotypes.

**Figure 2.**
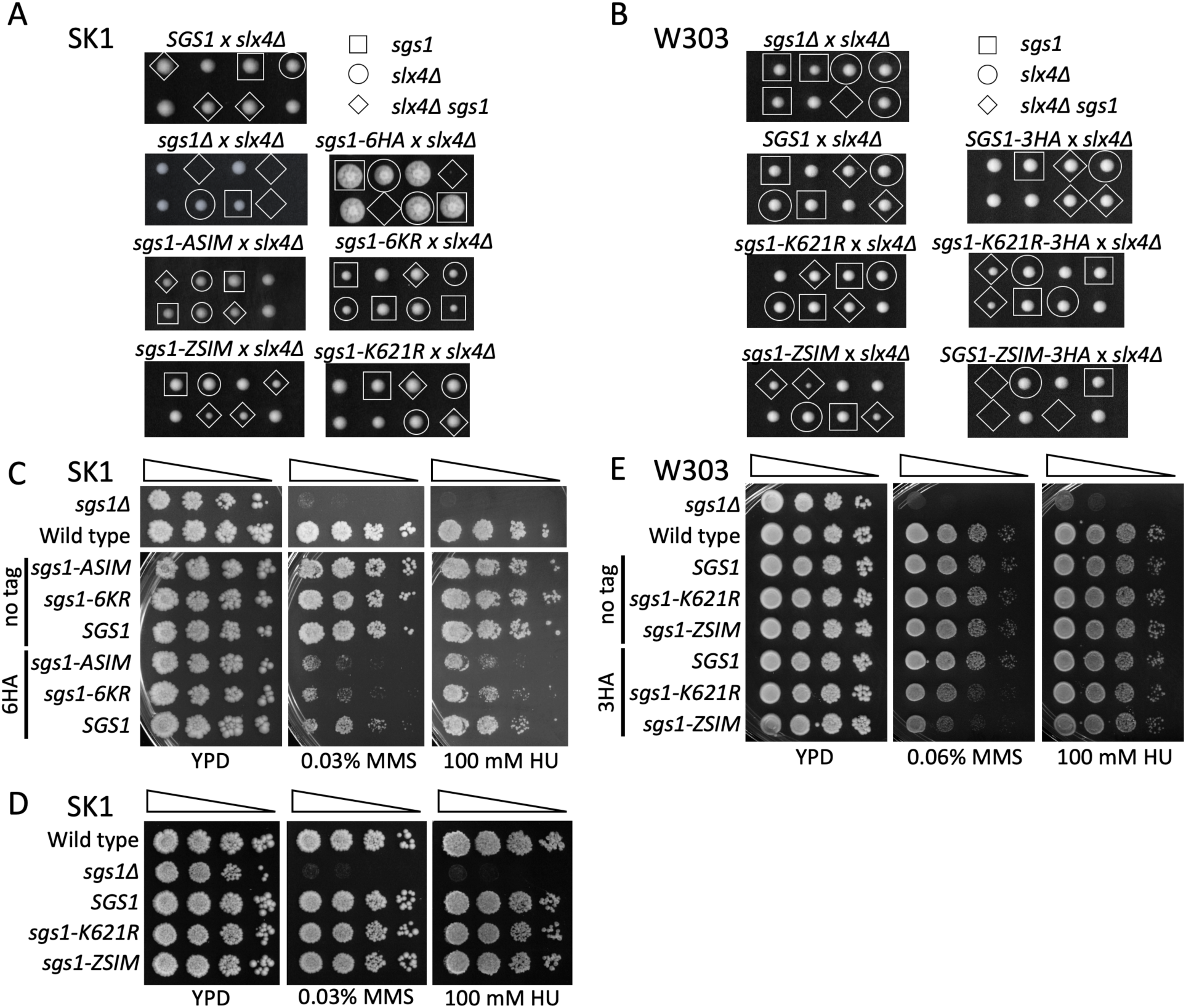
C-terminal HA tags exacerbate the mitotic phenotypes of *SGS1* SUMO-site mutants. In all panels, “*SGS1”* indicates an allelic replacement with wild-type sequences in the construct shown in Figure 1; “Wild type” indicates the unmodified endogenous *SGS1* locus. A) Genetic interactions between *slx4Δ* and *SGS1* SUMO-site mutants in SK1 strains. Two representative tetrads from the indicated heterozygous diploid are shown after grown for 2 days at 30°C. B) As in A, except with W303 strains from Bonner et al. (2016). C) MMS and HU sensitivity, in SK1, of *SGS1* SUMO-site mutants used by Bermudez-Lopez et al. (2016), with and without 6HA tags. 10-fold dilution series were spotted and allowed to grow 2-3 days at 30°C. D) As in C, but with SUMO-site mutants used by Bonner et al. (2016) without an epitope tag. E) As in D, but with tagged and untagged SUMO-site mutants in W303.

### Phenotypes of SGS1 mutants are exacerbated by a C-terminal HA tag

We first sought to reproduce some of the *SGS1* SUMO-site mutant phenotypes previously reported (Bermudez-lopez *et al*. 2016; Bonner *et al*. 2016). We found that the C-terminal 6HA tag used by Bermudez-Lopez *et al*. (Bermudez-Lopez *et al*. 2016) seriously compromised mitotic function of an otherwise wild-type *SGS1* gene in strains of the SK1 background. *sgs1-6HA* strains displayed synthetic lethality with *slx4Δ* (Fig. 2A), and also displayed MMS and HU sensitivity, although not to the same degree as an *sgs1Δ* mutant (Fig. 2C). Conversely, when present in an untagged context, *sgs1-ASIM* (which mutates SIMs in Sgs1) and *sgs1-6KR* (which lacks six lysines that are substrates for SUMOylation) did not show synthetic interaction with *slx4Δ* and were not sensitive to MMS or HU (Fig. 2A and 2C). However, the addition of a 6HA tag to these two mutants increased MMS and HU sensitivity relative to the tagged wild-type gene (Fig. 2C, Fig. S1). Thus, at least in the SK1 strain background, a C-terminal 6HA tag interferes with Sgs1 protein function and exacerbates SUMO-site mutant phenotypes.

We also found that the C-terminal 3HA tag used by Bonner *et al*. (Bonner *et al*. 2016) compromises Sgs1 function. Bonner *et al*. used W303 strains to examine mutants in the major Sgs1 SIM (*sgs1-ZSIM*, Fig. 1A) and a mutant lacking the major Sgs1 SUMOylation target lysine (*sgs1-K621R*, Fig. 1A), all in the context of an *SGS1* gene containing a C-terminal 3HA tag. They reported that *sgs-K621R-3HA* exhibits synthetic sickness with *slx4Δ*, and that *sgs1-ZSIM-3HA* is synthetically lethal with *slx4Δ*. We examined the same mutants in the absence of an epitope tag in the SK1 background and observed no synthetic interaction between *sgs1-K621R* and *slx4Δ*, and reduced growth, but not lethality, when *sgs1-ZSIM* is combined with *slx4Δ* (Fig. 2A). Furthermore, unlike the 3HA-tagged W303 mutants, the untagged mutants in SK1 displayed wild-type sensitivity to MMS and HU (Fig. 2D, Fig. S2A). These results further suggest that *SGS1* function is, at most, only partially affected by the *SGS1* SUMO-site mutants.

To test whether these phenotypic differences could be attributed to strain background differences, we removed the 3HA tag from the original W303 strains (Bonner *et al*. 2016) and restored the native *SGS1* terminator (Fig. 1C and Materials and Methods). When tested for synthetic interactions with *slx4Δ*, these untagged W303 strains behaved identically to the untagged SK1 strains, while 3HA-tagged strains recapitulated previously published phenotypes (Fig. 2B). In addition, the 3HA-tagged *SGS1* SUMO-site mutants, but not *SGS1-3HA*, displayed a modest increase in MMS sensitivity, while the untagged mutants were no more MMS-sensitive than wild type (Fig. 2E). These results indicate that both 3HA and 6HA C-terminal tags interfere with normal Sgs1 function, and that many of the phenotypes reported for *SGS1* SUMO-site mutants are due, in large part, to the presence of the epitope tag. We therefore performed all further analyses in SK1 strains lacking C-terminal tags.

### SGS1 SUMO-site mutants do not recapitulate the meiotic defects of mms21 SUMOylation-deficient or sgs1 loss-of-function mutants

Similar meiotic phenotypes are observed in *sgs1* loss-of-function mutants and SUMO ligase-defective *mms21-11* mutants, including reduced spore viability, bypass of *zmm* mutant phenotypes, and recombination and nuclear division defects when combined with mutants lacking the Mus81-Mms4 nuclease (Jessop and Lichten 2008; Oh *et al*. 2008; De Muyt *et al*. 2012; Zakharyevich *et al*. 2012; Xaver *et al*. 2013; Kaur *et al*. 2015; Tang *et al*. 2015). We first examined spore viability as a test of whether or not *SGS1* SUMO-site mutants show similar defects. As a control, we used *sgs1-md* (*pCLB2-SGS1*), in which Sgs1 is expressed during vegetative growth but is progressively depleted from the cells during meiosis (Lee and Amon 2003; Jessop and Lichten 2008; Oh *et al*. 2008). *sgs1-md* strains display a modest reduction in spore viability (90% compared to 98% in the wild-type and 97% in *SGS1-WT* allelic replacement, p<0.0001; ≥ 70 tetrads per genotype analyzed), similar to the defect observed in *mms21-11* mutants (89%; (Xaver *et al*. 2013). None of the untagged *SGS1* SUMO-site mutants showed reduced spore viability, while a strain with a 6HA-tagged wild-type *SGS1* (*sgs1-6HA*) showed reduced spore viability, to a greater extent than did *sgs1-md* (76% compared to 90% for *sgs1-md*, p<0.0001) (Fig. 3A). We do not understand the basis of the more severe phenotype of *sgs1-6HA*. It could be due to pre-meiotic defects that arise during vegetative growth; alternatively, it is possible that the 6HA tag creates an unproductive STR complex that sequesters Top3-Rmi1 and prevents it from performing its Sgs1-independent functions in joint molecule resolution (Kaur *et al*. 2015; Tang *et al*. 2015).

**Figure 3.**
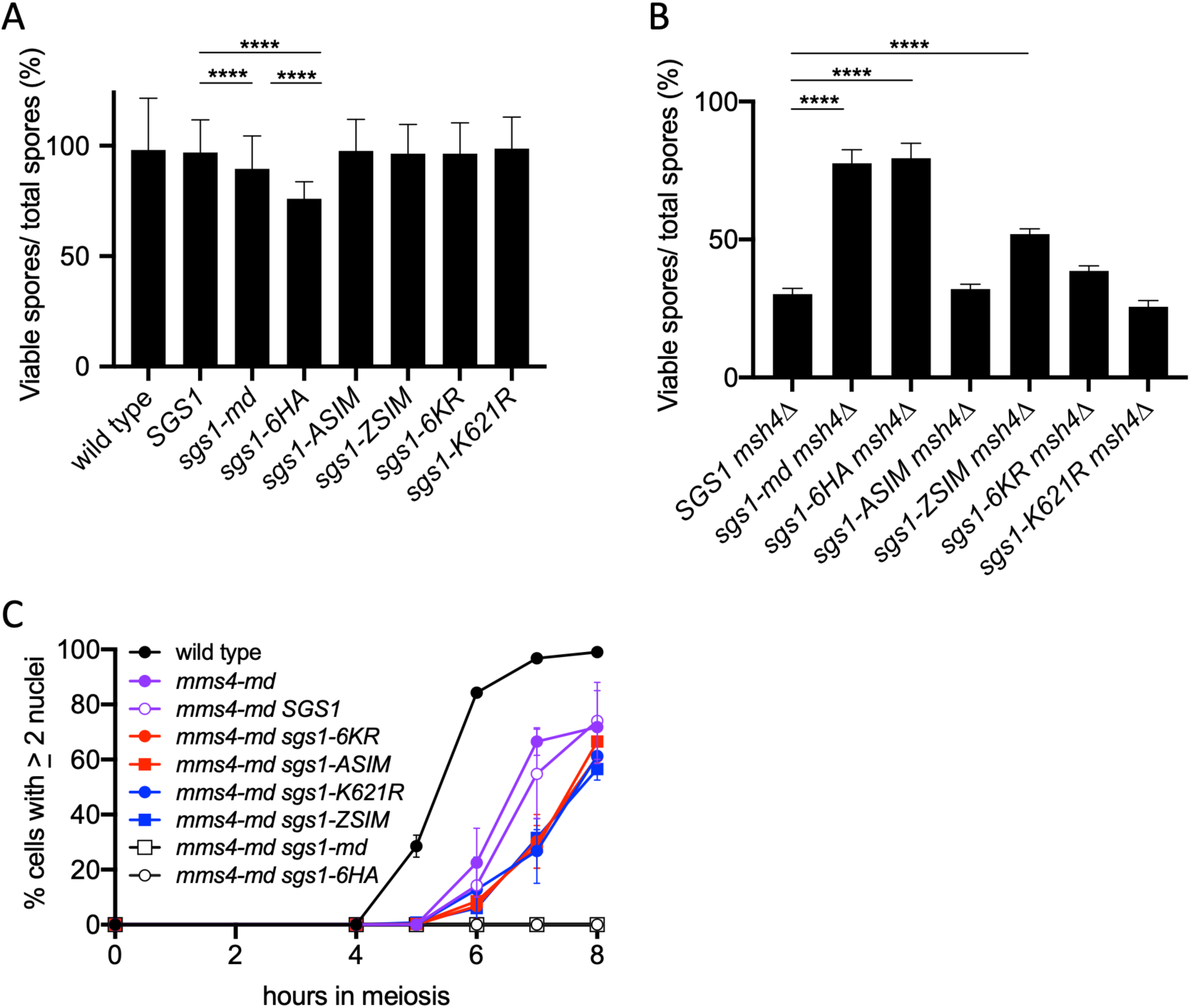
*SGS1* SUMO-site mutants retain STR meiotic function. In all panels, “wild type” denotes an unmodified *SGS1* locus; “*SGS1”* denotes the allelic replacement construct (see Figure 1) containing wild-type *SGS1* sequences. A) Spore viability of diploids homozygous for the indicated *SGS1* allele. Error bars represent the 95% confidence interval. **** indicates p<0.0001. B) Spore viability of diploids homozygous for *msh4Δ* and the indicated *SGS1* allele; other details as in A). C) Meiotic progression in diploids homozygous for the indicated genotype. The number of nuclei per cell was counted; cells with ≥ 2 nuclei were considered to have undergone at least one nuclear division. At least 200 cells were counted per time point in two independent experiments; error bars represent range.

*sgs1* and *mms21* loss-of-function mutants partially suppress the crossover defects and reduced spore viability phenotypes of *zmm* mutants, which have defects in the stabilization of regulated dHJ intermediates and their resolution as crossovers (Jessop *et al*. 2006; Lynn *et al*. 2007; Oh *et al*. 2007; De Muyt *et al*. 2012; Zakharyevich *et al*. 2012; Xaver *et al*. 2013). We asked if *SGS1* SUMO-site mutants suppress the spore inviability phenotype of strains lacking the ZMM protein Msh4 (Fig. 3B, Supplementary File 1). *sgs1-6HA* restored spore viability to *msh4Δ* to the same extent as *sgs1-md* (77% for *msh4Δ sgs1-md*, 80% for *msh4Δ sgs1-6HA*, compared to 30% for *msh4Δ SGS1-WT*). In contrast, most untagged *SGS1* SUMO-site mutants did not significantly increase *msh4Δ* spore viability. The only exception was *sgs1-ZSIM*, which showed partial suppression of the *msh4Δ* spore viability defect, but not to the same extent as *sgs1-md* (52% for *msh4Δ sgs1-ZSIM*, 77% for *msh4Δ sgs1-md*). The partial suppression seen with untagged *sgs1-ZSIM* is consistent with the synthetic slow growth phenotype of *sgs1-ZSIM slx4Δ* double mutants, and suggests that this mutation confers a greater loss-of-function than the other *SGS1* SUMO-site mutants.

To more stringently test meiotic Sgs1 function, we examined genetic interactions between *SGS1* SUMO-site mutants and *mms4-md* (*pCLB2-MMS4*), which does not express Mms4 during meiosis (De Muyt *et al*. 2012). When STR activity is absent, efficient meiotic recombination intermediate resolution requires Mus81-Mms4. Importantly, *sgs1 mus81* and *sgs1 mms4-md* double mutants display nuclear division failure caused by unresolved recombination intermediates (Jessop and Lichten 2008; Oh *et al*. 2008; De Muyt *et al*. 2012; Zakharyevich *et al*. 2012). The same phenotype is observed in *mms4-md mms21-11* double mutants (Xaver *et al*. 2013), so we asked if any of the *SGS1* SUMO-site mutants recapitulate this synthetic interaction (Fig. 3C). As expected, *sgs1-md mms4-md* double mutants displayed a complete block to nuclear divisions, as did *sgs1-6HA mms4-md* double mutants, consistent with the C-terminal HA tag disrupting *SGS1* meiotic function. *mms4-md* single mutants displayed a slight delay in nuclear division relative to wild-type, and strains where *mms4-md* was combined with any of the untagged *SGS1* SUMO-site mutants displayed a further 1-hour delay (Fig. 3C). The delayed nuclear divisions in *mms4-md SGS1* SUMO-site double mutants suggests that an increased fraction of recombination intermediates utilize the non-ZMM resolution pathway in these strains, consistent with the SUMO-site mutants conferring a partial loss of *SGS1* meiotic function.

## Discussion

This report shows that the presence of a C-terminal HA tag is largely responsible for the previously reported phenotypes of *SGS1* mutants proposed to influence Mms21-mediated modification of the STR complex (Bermudez-Lopez *et al*. 2016; Bonner *et al*. 2016). We found that both 3HA and 6HA C-terminal tags on Sgs1 sensitize cells to MMS and HU and exacerbate SUMO-site mutant phenotypes; this effect was observed in two different *S. cerevisiae* genetic backgrounds. It is likely that these effects were missed in previous studies because tagged and untagged mutant alleles were not compared. For example, in W303 strains, an *SGS1-3HA* construct did not display a synthetic interaction with *slx4Δ* or increased sensitivity to either MMS or HU (Bonner *et al*. 2016). However, we found that the presence of a 3HA tag causes, for both *sgs1-K621R-3HA* and *sgs1-ZSIM-3HA*, a more severe synthetic interaction with *slx4Δ* and increased MMS sensitivity than when the tag is absent (Fig. 2). We also find that, in SK1 strains, a C-terminal 6HA tag on an otherwise wild-type *SGS1* causes complete synthetic lethality with *slx4Δ* and sensitivity to both MMS and HU (Fig. 2); this allele also causes reduced spore viability, suppresses the reduced spore viability of *msh4Δ*, and blocks meiotic nuclear division when combined with *mms4-md* (Fig. 3). These phenotypes are suggestive of substantial loss of Sgs1 function. It therefore appears that a C-terminal HA tag can compromise Sgs1 function, but this compromised function may not be detected until the HA tag is combined with other mutants or until the strain is sufficiently challenged by other means.

While we do not know the exact nature of the defects caused by C-terminal HA tags, our findings suggest that the Sgs1 C-terminus may have an important biological function. The Sgs1 C-terminus contains a region predicted to be highly unstructured. This region follows immediately after the conserved HRDC (Helicase-and-RNaseD-like-C-terminal) domain, which is important for substrate binding and regulation of helicase activity (Liu *et al*. 1999; Yankiwski *et al*. 2001; Hickson 2003; Wu *et al*. 2005; Chu and Hickson 2009; Vindigni and Hickson 2009; Harami *et al*. 2017). The HA epitope has a net negative charge, and could conceivably interfere with Sgs1-DNA interaction, especially in the HRDC domain. Thus, it is possible that the tag interferes with the biochemical function of this HRDC domain and/or the disordered C-terminus, thereby compromising Sgs1 function during D-loop disruption or dHJ dissolution. This would result in the persistence of DNA structures whose resolution requires endonucleases such as Mus81-Mms4 and Slx1-Slx4, which in turn could account for the synthetic lethality observed in *sgs1-6HA slx4Δ* strains and the meiotic nuclear division failure observed in *sgs1-6HA mms4-md* diploids.

The absence of strong mitotic or meiotic defects in untagged *SGS1* SUMO-site mutants prompted us to examine genetic interactions of these alleles with mutants in genes known to confer synthetic phenotypes when combined with *sgs1Δ* and *mms21-11*. While *sgs1* loss of function and *mms21-11* mutants strongly suppress the spore inviability of *zmm* mutants (De Muyt *et al*. 2012; Xaver *et al*. 2013; Tang *et al*. 2015; Fig. 3B), of the four SUMO-site mutants examined, only *sgs1-ZSIM* significantly suppressed *msh4Δ*, and this suppression was partial (Fig. 3B). In a similar vein, while meiotic Sgs1 depletion (in *sgs1-md*) led to a complete block to nuclear division in *mms4-mn* mutants, the *SGS1* SUMO-site mutants delayed division by 1h (Fig. 3C), suggesting that these mutants confer, at most, a partial defect in STR complex activity.

The suggestion that SUMO-site mutants only slightly impinge upon STR complex function could be explained in a number of ways. For example, if the SUMO-site mutants studied do substantially reduce STR complex SUMOylation, as was suggested in previous studies (Bermudez-Lopez *et al*. 2016; Bonner *et al*. 2016), it is possible STR activity is regulated by factors or post-translational modifications other than Mms21-mediated SUMOylation. Alternatively, it is possible that, in the absence of epitope tags, the SUMO-site mutants we studied only partially disrupt Mms21-mediated SUMOylation of the STR complex, either because alternative SUMOylation sites (Klug *et al*. 2013; Pichler *et al*. 2017) or because other SUMO ligases (Klug *et al*. 2013; Hendriks and Vertegaal 2016; Pichler *et al*. 2017) come into use. Testing these possibilities will require isolation of antibodies to the native Sgs1 protein, or identification of epitope tags that do not disrupt Sgs1 function. Finally, since *SGS1* SUMO-site mutants did not recapitulate, in any of the assays used, the phenotypes of SUMO E3 ligase-null *mms21-11* mutants, it is possible that Mms21 modification of proteins other than the STR complex is required for proper recombination and repair during mitosis and meiosis. In this regard, it will be of interest to determine the full set of factors that Mms21 acts through to exert its functions. We note that a recent analysis of the yeast meiotic SUMO proteome found SUMO conjugated to many recombination-associated proteins, including several subunits of the Smc5-Sm6-Mms21 complex (Bhagwat *et al*. 2019).

A recent report (Zapatka *et al*. 2019) has proposed that Mms21-mediated SUMOylation of Smc5 is involved in error-free bypass of damaged replication forks. It is worth noting that, in this study, mutations in *SMC5* thought to interfere with its SUMOylation did not have strong phenotypes, but exhibited synthetic chromosome segregation defects when combined with *mms4Δ*. This finding is reminiscent of our finding that meiotic nuclear divisions are delayed in *mms4-md SGS1* SUMO-site double mutants. (Fig. 3C).

In summary, our data show that the phenotypes of mutants thought to affect the STR interaction with and modification by the Smc5-Smc6-Mms21 complex are exacerbated by a C-terminal tag on *SGS1* and reveal a possible role for the C-terminus of Sgs1 in recombination. These finding reinforce the importance of testing genetic interactions of mutants without a protein tag to ensure that the effects seen are not due to an exacerbating effect caused by the tag, but rather represent true biological function.

## Acknowledgements

We thank Xiaolan Zhao and Luis Aragon for strains and plasmids, Xiaolan Zhao and Martin Xaver for helpful discussions and technical advice, and Jasvinder Ahuja, Matthew Bochman, Alexander Kelly, Soni Lacefield, Hengyao Niu, and Martin Xaver for comments on the manuscript. This work was supported by the Intramural Research Program of the NIH through the Center for Cancer Research at the National Cancer Institute.

---

**Figure S1.**
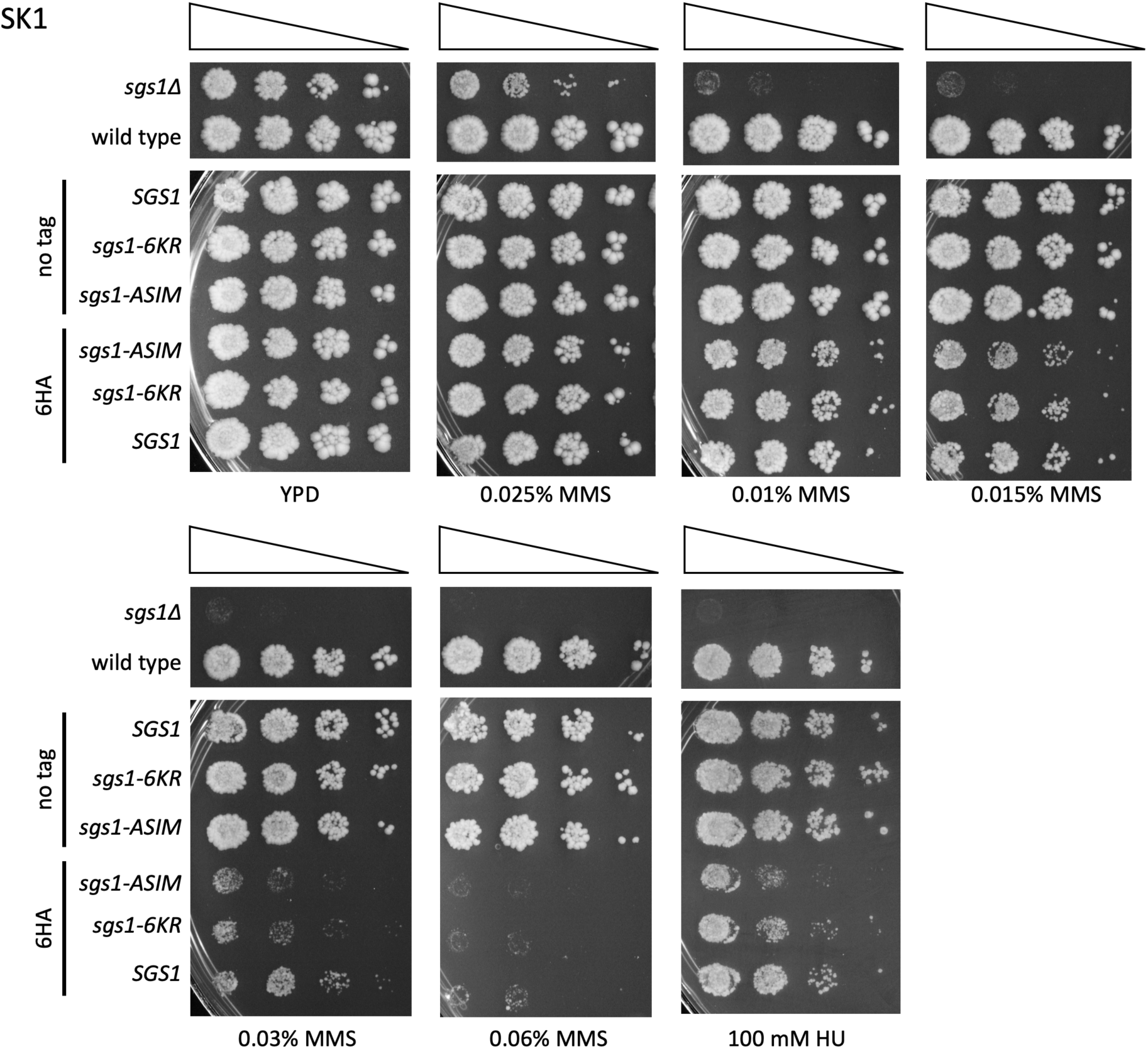
MMS-sensitivity of *SGS1* SUMO-site mutant alleles from Bermudez-Lopez et al (2016). SK1 strains with the indicated *SGS1* alleles (see Figures 1 and 2 for details) were spotted in a 10-fold dilution series on plates with different MMS concentrations and incubated at 30°C for 2-3 days. Note that SUMO-site mutants without a 6HA tag show wild-type resistance to MMS and HU, while, in the context of a 6HA tag, SUMO-site mutants are slightly more sensitive than the already-sensitive 6HA-tagged wild-type allele.

**Figure S2.**
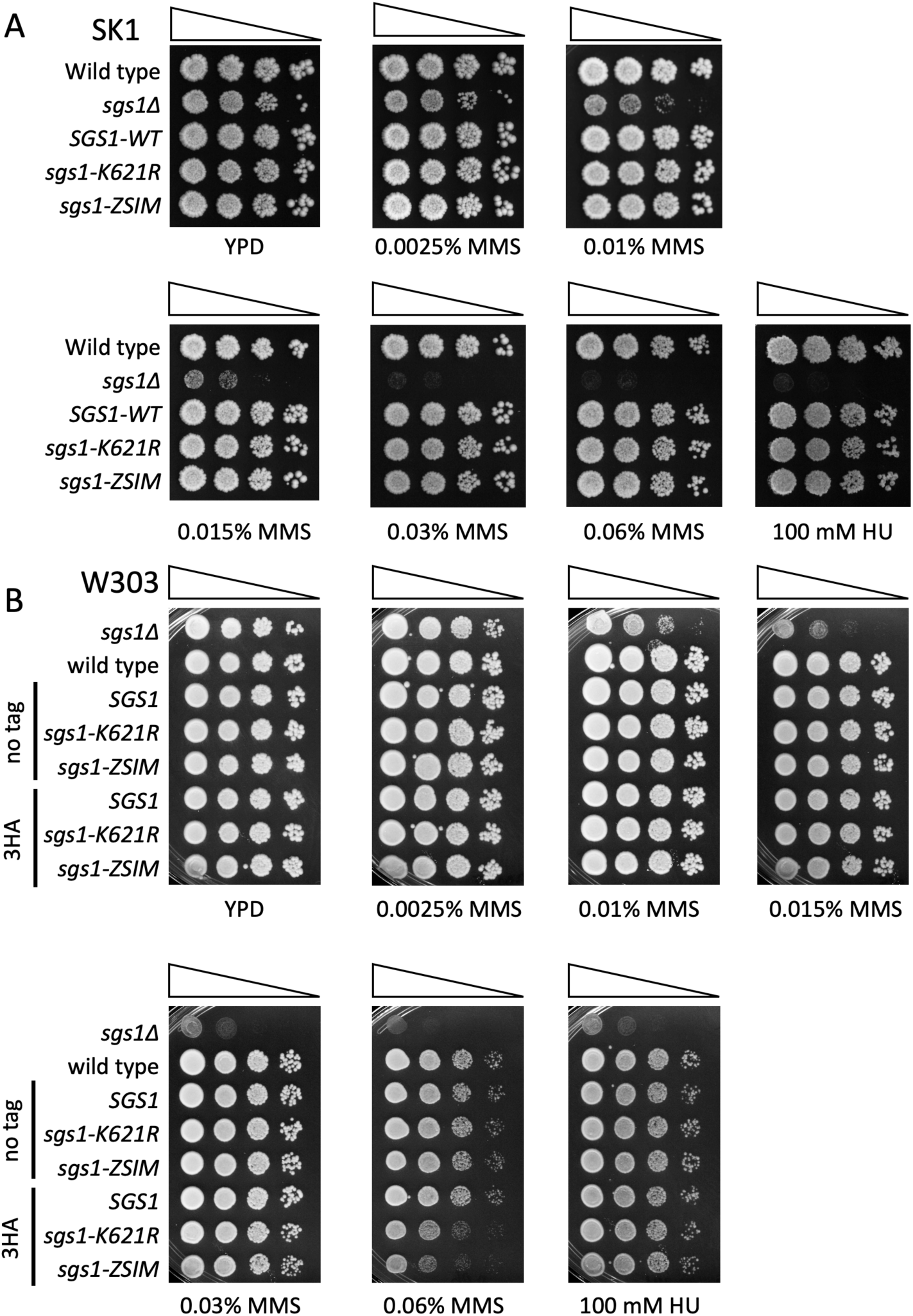
MMS sensitivity of *SGS1* SUMO-site mutant alleles from Bonner et al. (2016). A) SK1 strains with the indicated *SGS1* alleles (all without a C-terminal tag) were spotted in a 10-fold dilution series on plates with different MMS concentrations and incubated at 30°C for 2-3 days. B) As in A, but with W303 strains. Note that in an untagged context, SUMO-site mutants show wild-type resistance to MMS, as does a strain with a 3HA-tagged *SGS1* allele. In the context of a 3HA C-terminal tag, SUMO-site mutants are slightly more sensitive to 0.06% MMS than is wild type.

**Table S1.**
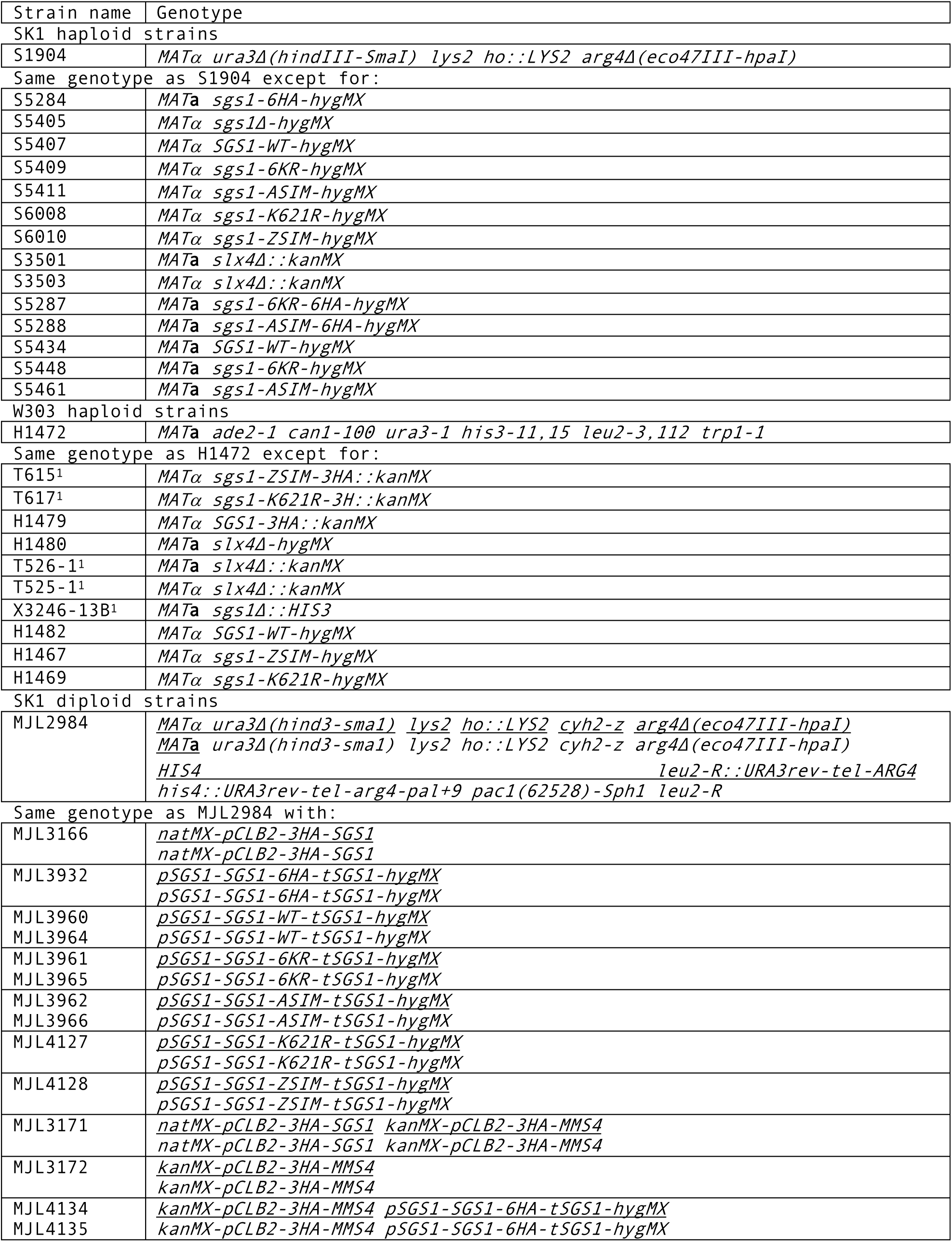
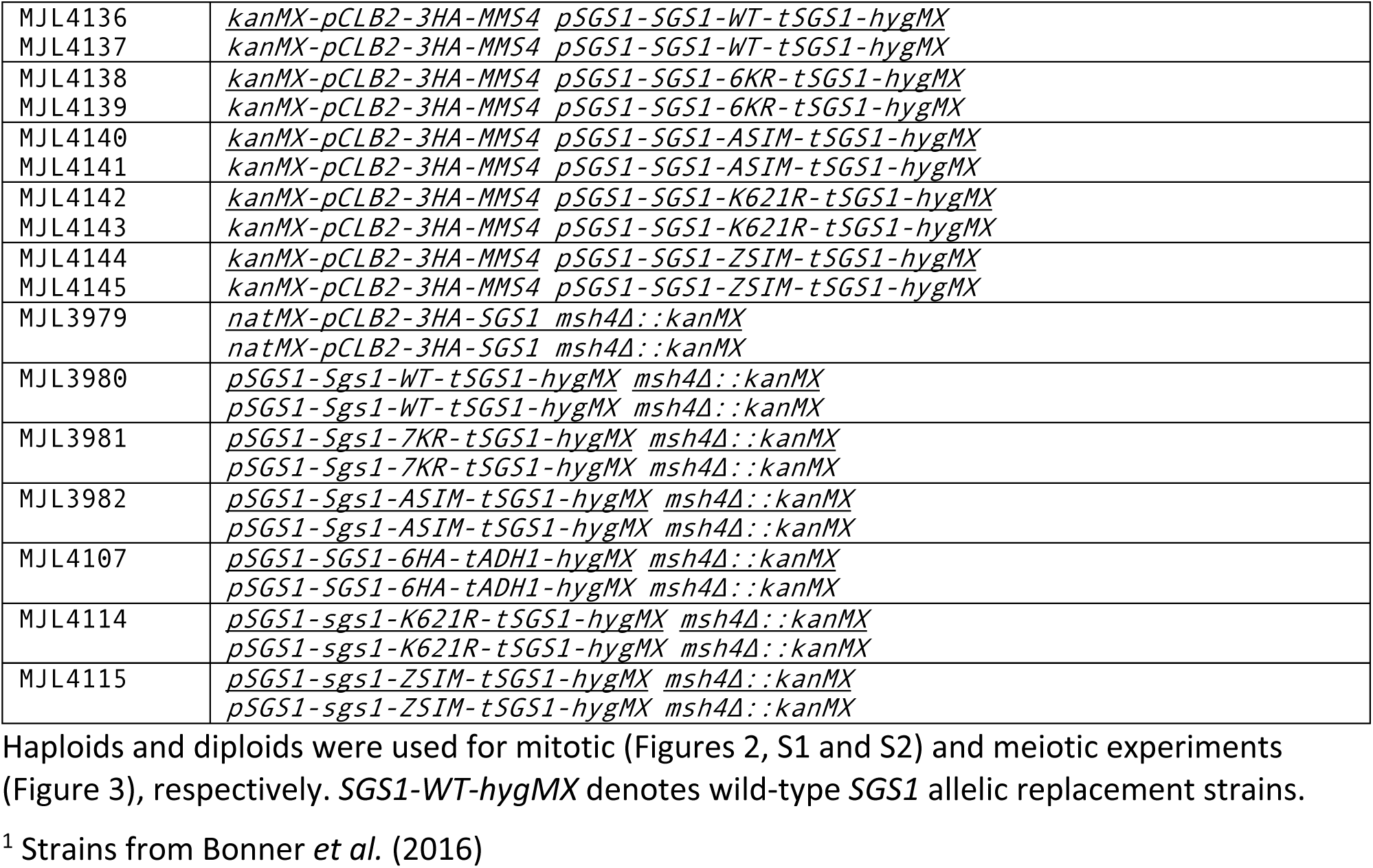
Strains used in this study.

